# Nicotine-mediated rescue of α-synuclein toxicity requires synaptic vesicle glycoprotein 2

**DOI:** 10.1101/2022.07.26.501592

**Authors:** Abby L. Olsen, Sabrina G. Clemens, Mel B. Feany

**Affiliations:** Brigham and Women’s Hospital, Department of Neurology, Chevy Chase, MD, 20815; Brigham and Women’s Hospital, Department of Pathology, Chevy Chase, MD, 20815; Aligning Science Across Parkinson’s (ASAP) Collaborative Research Network, Chevy Chase, MD, 20815

## Abstract

**Background:** Parkinson’s disease (PD) is characterized by α-synuclein aggregation and loss of dopamine (DA) neurons in the substantia nigra. Risk of PD arises due to a combination of genetic and environmental factors, which may interact, termed gene-environment (GxE) interactions. An inverse association between smoking and risk of PD is well-established, and a previous genome-wide GxE interaction study identified genetic variation in the synaptic-vesicle glycoprotein 2C (*SV2C*) locus as an important mediator of the degree to which smoking is inversely associated with PD.

**Objective:** We sought to determine the mechanism of the smoking-*SV2C* interaction in a *Drosophila* model of PD.

**Methods:** Flies expressing human α-synuclein in all neurons develop the hallmarks of PD, including motor dysfunction, loss of DA neurons, and formation of α-synuclein inclusions. We assessed the effects of increasing doses of nicotine on these parameters of neurodegeneration, in the presence or absence of *SV2* knockdown.

**Results:** α-synuclein-expressing flies treated with nicotine had improvement in locomotion, DA neuron counts, and in α-synuclein aggregation. However, in α-synuclein-expressing flies in which Drosophila orthologs of *SV2* were knocked down, nicotine failed to rescue neurodegeneration.

**Conclusions:** This work confirms a GxE interaction between nicotine and *SV2*, defines a role for this interaction in α-synuclein proteostasis, and suggests that future clinical trials on nicotine should consider genetic variation in *SV2C*. Further, this provides proof of concept that our model can be used for mechanistic study of GxE, paving the way for investigation of additional GxE interactions or identification of novel GxE.

## Introduction

Parkinson’s disease (PD) is the second most common neurodegenerative disease, affecting 1% of the population over the age of 60. It arises due to a combination of genetic and environmental factors. How genes and environment interact together to influence risk remains incompletely understood. Traditional genome-wide association studies (GWAS) have been effective at identifying associated risk alleles, such as *SNCA, LRRK2*, and *GBA*, but can miss interactions with low effect sizes. Indeed, even the largest GWAS to date only accounts for a minority of heritable risk of PD^1^. Similarly, epidemiologic studies have identified numerous environmental factors that are positively or negatively associated with PD, but these studies can be challenging to reproduce, and causality difficult to determine. Thus, there is a need for mechanistic studies investigating gene-environment interactions (GxE).

By stratifying subjects into groups based on environmental exposures and performing a genome-wide association and interaction study (GWAIS), genes that exert their effects only in the context of certain environmental exposures can cross the threshold of significance. In a previous GWAIS, we identified an interaction between smoking and the synaptic vesicle protein *SV2C* in PD patients^2^. That is, the degree to which smoking was associated with PD was dependent on genetic variation in the *SV2C* locus^2^, ranging from strongly negatively associated to strongly positively associated, with an odds ratio (OR) of 0.44 – 3.23, depending on genotype at two independent SNPs.

SV2C belongs to a family of synaptic vesicle glycoproteins, along with SV2A and SV2B. The SV2 family proteins are transmembrane proteins found on secretory vesicles, including synaptic vesicles, and they have been reported to facilitate exocytosis of synaptic vesicles by rendering them responsive to calcium^3,4^. Expression of *SV2C* is primarily localized to the basal ganglia, with highest levels found in dopaminergic (DA) neurons of the substantia nigra (SNpc), ventral tegmental area (VTA), and in cholinergic interneurons of the striatum^5^. Certain *SV2C* variants in PD patients have been shown to predict sensitivity to levodopa^6^, and *SV2C* expression pattern is altered in PD patients^7^. Thus, *SV2C* and other SV2 family members represent credible targets as potential modifiers of PD.

Similarly, smoking has been inversely associated with PD in numerous epidemiologic studies. Although there are different hypotheses proposed to explain this relationship, including decreased sensation-seeking behavior in PD^8^, most evidence suggests that nicotine contributes significantly to neuroprotection against PD^9^. Nicotine has been shown to stimulate DA release in smokers^10^ and increase degradation of SIRT6, a pro-inflammatory protein^11^. Further, cholinergic deficits are also present in PD^12^, which could conceivably be alleviated by nicotine as a nicotinic acetylcholine receptor (nAChR) agonist. There is evidence for a protective effect of nicotine *in vivo*, with animal models of PD showing amelioration after nicotine exposure^13,14^. Results of nicotine in clinical trials have been mixed, however^15–23^. Further, even in trials demonstrating some benefit of nicotine, it is unknown whether this effect is merely symptomatic, alleviating symptoms, or neuroprotective, decreasing disease progression. Finally, it is possible that nicotine-free tobacco may also confer neuroprotection, suggesting a role for non-nicotinic mechanisms^24^. Thus, while nicotine is the most biologically plausible factor underlying the inverse relationship between smoking and PD, many questions regarding its mechanism remain.

In our prior work, in addition to identifying the smoking-*SV2C* interaction in PD patients, we also identified a gene-environment interaction between nicotine and a *Drosophila* ortholog of the *SV2* family, in a paraquat-induced model of parkinsonism^2^. Here we sought to validate this interaction in an α-synuclein based *Drosophila* model of PD and to determine the mechanism underlying the interaction.

## Methods

### Drosophila genetics

All fly crosses and aging were performed at 25 °C. All experiments were performed at 10 days post-eclosion unless otherwise noted in the figure legends. All experiments include roughly equal numbers of male and female flies in which wild type human α-synuclein is expressed in neurons using the bipartite QUAS-QF2 expression system^25^ and the pan-neuronal driver *neuronal-synaptobrevin (nSyb)-QF2*. Additionally, genes of interest were knocked down using the UAS-GAL4 system, also driven by *n-synaptobrevin*. The *UAS-Atg8a-GFP* and *UAS-GFP-mCherry-Atg8a* stock and the transgenic RNAi stocks *UAS-CG3168 JF02441* and *UAS-CG14691 JF03420* were obtained from the Bloomington *Drosophila* Stock Center. The transgenic RNAi stocks *UAS-CG3168 48010* and *UAS-CG14691 46365* were obtained from the Vienna *Drosophila* Resource Center. Flies lacking α-synuclein or the RNAi but containing both drivers were used as controls. The genotype for α-synuclein expressing flies was *QUAS-Syn, nSyb-QF2, nSyb-GAL4/+* or *QUAS-Syn, nSyb-QF2, nSyb-GAL4/UAS-RNAi*, unless otherwise noted in the figure legend. These flies were compared to control flies expressing the same genetic drivers but lacking α-synuclein, that is, *nSyb-QF2, nSyb-GAL4/+* or *nSyb-QF2, nSyb-GAL4/UAS-RNAi*.

### Drug feeding and maintenance

*Drosophila* crosses were performed on standard fly food. At the time of eclosion flies were placed in vials containing a 1:1 ratio of instant medium (Carolina) to aqueous nicotine solution. Nicotine (Sigma) was diluted in 100% ethanol and then further diluted with distilled water to a final concentration of 0.1, 0.2, or 0.4 mg/ml in 4% ethanol. Control flies grown in the absence of drug were fed instant medium mixed with vehicle control (4% ethanol).

### qRT-PCR

To confirm the knock down of selected genes, exon-spanning primers were selected from the DRSC FlyPrimerBank. 10 fly heads per condition were homogenized in Qiazol (Qiagen) and total RNA was isolated. Samples were treated with DNAse and cDNA prepared using a High Capacity cDNA Reverse Transcription Kit (Applied Biosystems). Amplification was reported by SYBR green in a QuantStudio 6 Flex System (ThermoFisher), and relative expression was determined using ΔΔCt method normalized to the *RPL32* housekeeping gene.

### Locomotion assay

Flies were sorted into cohorts containing 9-14 flies per vial. At day 7 of exposure to either vehicle or drug, flies were transferred to a clean vial (without food) and allowed to acclimate for 1 min. The locomotion assay was performed as previously described^26^. Six technical replicates were averaged for each biological replicate, and there were six biological replicates for each condition. Significant difference between phenotypes at the same time-point was measured by two-way ANOVA with Tukey’s multiple comparison test. Two RNAi lines per gene were used for behavior studies (Figure 1, Supplemental Figure 1), and a single RNAi was used for all subsequent confirmatory pathologic studies. The RNAi lines used in the confirmatory studies were the *UAS-CG3168 JF02441* and *UAS-CG14691 JF03420* lines. After identifying a dose response in locomotion to nicotine of 0.1-0.4 mg/ml (Figure 1), the highest dose of 0.4 mg/ml was used for all subsequent confirmatory pathologic studies.

**Figure 1:**
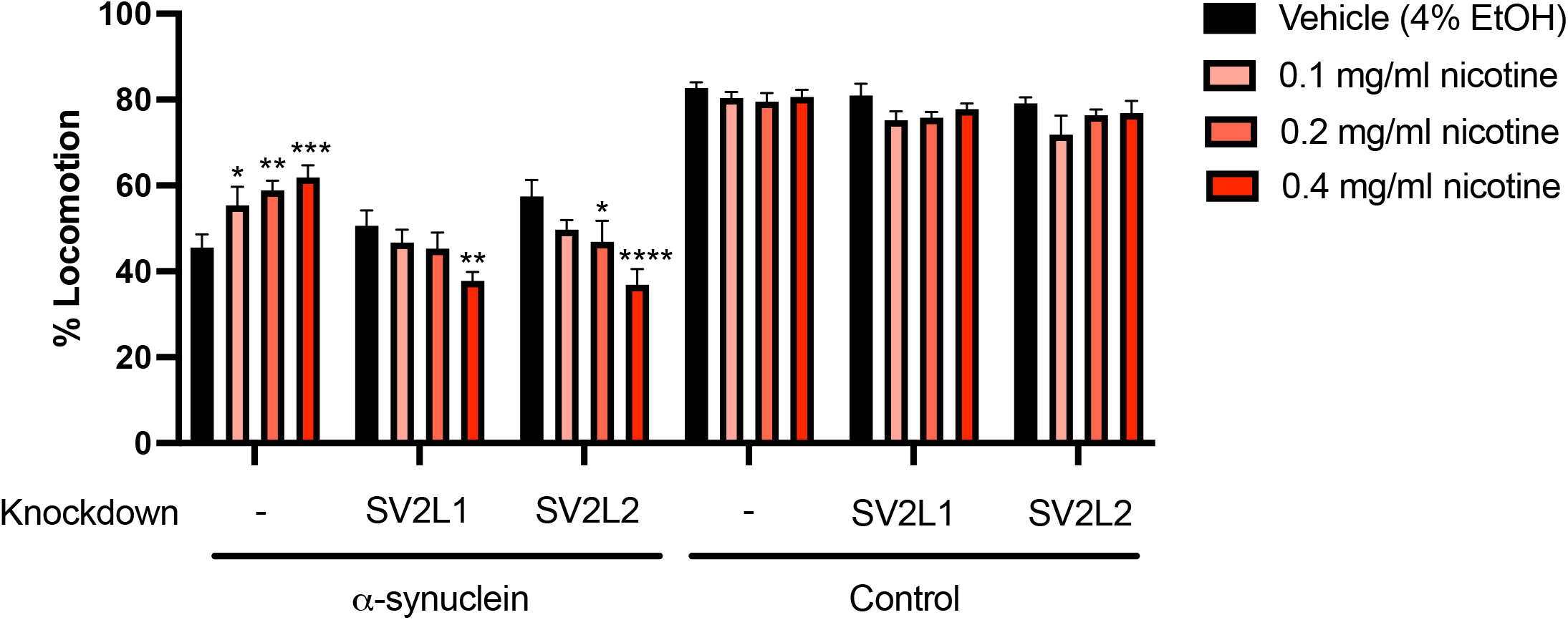
Rescue of α-synuclein induced locomotion impairment by nicotine requires *SV2*. α-synuclein or control flies were fed media with 0.1-0.4 mg/ml nicotine or vehicle control from the time of eclosion up to 10 days of age. Locomotion was measured at 4, 7, and 10 days. The day 7 timepoint is shown; results at 4 and 10 days were similar. Nicotine improved locomotion in α-synuclein expressing flies (far left) in a dose-dependent manner. However, nicotine failed to rescue the locomotor deficit when the *SV2* orthologs *SV2L1* and *SV2L2* were knocked down and in fact further worsened locomotion in this condition in a dose-dependent manner. N = minimum 6 biological replicates of 9-14 flies each per genotype. *p <0.05, **p < 0.01, *** p < 0.005, **** p < 0.001. Within each genotype, statistical significance was determined using two-way ANOVA with Dunnet’s multiple comparison test, comparing each dose of nicotine to vehicle control.

### Immunohistochemistry & Immunofluorescence

After 10 days of exposure to either vehicle or 0.4 mg/ml nicotine, flies were anesthetized, fixed in formalin overnight, and embedded in paraffin wax. A series of frontal sections of fly brain were cut either at 2 or 4 μm in thickness and processed through xylenes, ethanol, and water. For total neuron counts (2 μm), slides were stained with hematoxylin. For DA neuron counts (4 μm), immunohistochemistry was performed. Antigen retrieval for 30 min in 10 mM sodium citrate, pH 6.0, was performed in a pressure cooker. After cooling and depressurization, slides were blocked in 2% milk hydrated in PBS with 0.3% Triton for 1 hr and then incubated with primary diluted in the same blocking solution at room temperature overnight. Antibodies used include Atg8a/LC3 (clone EIJ4E, 1:2,000, rabbit, Cell Signaling), and ref(2)P/p62 (1:5,000 rabbit). The next day slides were washed and incubated for 1 hr in a secondary biotin-conjugated anti-mouse antibody (1:200, goat anti mouse, SouthernBiotech, Birmingham, AL) mixed in blocking solution. The ubiquitin antibody is directly conjugated to horse radish peroxidase and therefore does not require a secondary antibody. Following washing, avidin–biotin–peroxidase complex (Vectastain Elite) mixed in PBS was applied for 1 hr followed by development using diaminobenzidine (ImmPACT DAB, Vector, Torrance, CA).

For immunofluorescence, microwave antigen retrieval was conducted in 10 mM sodium citrate, and slides were blocked in 2% milk hydrated in PBS with 0.3% Triton for 1 hr. All primary antibodies were diluted in blocking solution. Primary antibodies used include tyrosine hydroxylase (1:200, mouse, Immunostar), α-synuclein 5G4 (1:50,000, mouse, Millipore), and GFP (1:5, Developmental Studies Hybridoma Bank). Slides were incubated with fluorophore-conjugated secondary antibodies in blocking solution for 1 hr (1:200, Alexa 488 or 555, Invitrogen) then mounted with DAPI-containing Fluoromount medium (SouthernBiotech). Tissue was excited and photographed using Zeiss LSM 800 confocal microscopy and analyzed with ImageJ.

### Quantification of neuron counts

Images of the anterior medulla from hematoxylin-stained formalin-fixed paraffin-embedded tissue were captured at 40X magnification using brightfield microscopy. One image per fly and 6 flies per genotype were used for quantification. The number of nuclei in each tissue section was counted and normalized to the area of the section.

### Quantification of TH+ neuron counts and α-synuclein aggregates

Images of the anterior medulla from immunofluorescence-stained formalin-fixed paraffin-embedded tissue were captured on a confocal microscope at 63X magnification. One image per fly and 6 flies per genotype were used for quantification. The number of TH+ cells or α-synuclein aggregates in each tissue section was counted and normalized to the area of the section.

### α-synuclein monomer detection

At day 1 post-eclosion, one fly head per condition was homogenized in 2X Laemmli buffer, boiled for 10 min, and centrifuged. SDS-PAGE was performed (Lonza) followed by transfer to PVDF membrane (Bio-Rad) and microwave antigen retrieval in PBS. Membranes were blocked in 2% milk in PBS with 0.05% Tween 20 for 1 hour, then immunoblotted with primary antibody α-synuclein H3C (1:100,000, mouse, Developmental Studies Hybridoma Bank) in 2% milk in PBS with 0.05% Tween-20 overnight at 4 °C. Membranes were incubated with appropriate horseradish peroxidase-conjugated secondary antibodies (1:50,000) in 2% milk in PBS with 0.05% Tween 20 for 3 hours. Signal was developed with enhanced chemiluminescence (Thermo Scientific).

### α-synuclein oligomer detection

20 fly heads per genotype were homogenized in 20 ul TNE lysis buffer (10 mm Tris HCl, 150 mM NaCl, 5 mM EGTA, 0.5% NP40) supplemented with HALT protease and phosphatase inhibitor (Roche). The homogenate was briefly spun down to remove debris. The remaining supernatant was ultracentrifuged at 100,000 x g for 1 hour at 4 C. The supernatant was transferred to a new tube and combined with 2X Laemelli buffer at a 1:1 ratio. SDS-PAGE was then performed as above but without microwave antigen retrieval. Prior to incubating with primary antibody, the blot was cut to separate monomeric from oligomeric α-synuclein. Primary antibody was α-synuclein clone 42 (1:5,000 mouse, BD Bioscience).

### Quantification of Atg8a and ref(2)P puncta

Images of the anterior medulla from immunohistochemistry-stained formalin-fixed paraffin-embedded tissue were captured at 40X magnification using brightfield microscopy. One image per fly and 6 flies per genotype were used for quantification. The number of large puncta (>0.25 μm^2^, measured in ImageJ using the particle analysis function) in each tissue section were counted and normalized to the area of the section.

### Measurement of autophagic flux

Freshly dissected brains from 10-day-old animals were imaged at 63X on a Zeiss LSM 800 confocal microscopy. GFP-mCherry-Atg8a structures were analyzed as previously described^27^. For analysis of mCherry-GFP ratio across puncta, GFP-mCherry-Atg8a puncta within the field of view were automatically selected using h_watershed in Fiji^28^. For each punctum, intensity of fluorescence was measured for each channel, and the ratio of red to green fluorescence was calculated.

#### Data Sharing

The data that support the findings of this study are available from the corresponding author, upon reasonable request.

## Results

### Nicotine rescues α-synuclein induced impaired locomotion in a dose-dependent manner

We have previously demonstrated that nicotine treatment improves lifespan in a dosedependent manner in paraquat-induced parkinsonism in the fly^2^. To determine if nicotine has neuroprotective effects in an α-synuclein based model, we first measured the effects on locomotion using a previously published assay^26^. We assessed locomotion at day 7 post-eclosion and determined that nicotine rescues impaired locomotion in a dose-dependent manner (Figure 1).

This rescue in locomotor activity due to nicotine is dependent on *SV2C. Drosophila* have multiple *SV2* family orthologs. We chose to examine two: *CG3168*, which is the most closely related to the human *SV2* family as determined by *Drosophila* RNAi Screening Center Integrative Ortholog Predictor Tool (DIOPT), and *CG14691*, which was identified in our prior work as interacting with nicotine^2^. Because of their homology with the larger synaptic vesicle glycoprotein 2 (*SV2*) gene family, we have name these *SV2L1* and *SV2L2*, respectively, for synaptic vesicle glycoprotein 2 like.

When *SV2L1* or *SV2L2* are knocked down (Figure 1), nicotine no longer increases locomotion, and in fact, the highest dose of nicotine (0.4 mg/ml) significantly worsens locomotion in the α-synuclein flies. This dose of nicotine was used for all subsequent experiments. We confirmed the locomotion result with a second RNAi line (Supplemental Figure 1). Effective knockdown of *SV2L1* and *SV2L2* with each RNAi is demonstrated in Supplemental Figure 2. All subsequent experiments use RNA #1 for each gene.

### Rescue of neurodegeneration by nicotine requires SV2

Compared to control flies, flies expressing α-synuclein in neurons demonstrate significant neurodegeneration, quantified by fewer cells per square micron in the densely cell-populated anterior medulla (Figure 2). When α-synuclein flies were exposed to 0.4 mg/ml of nicotine, we observed a rescue of cell density. The effect of nicotine was abrogated with knockdown of *SV2L1* and *SV2L2*. Similarly, nicotine rescued loss of dopaminergic neurons (Figure 3), which also required *SV2L1* and *SV2L2* expression. Interestingly, with *SV2L1* and *SV2L2* knockdown, nicotine not only failed to rescue dopaminergic neuron loss but in fact further significantly exacerbated it, similar to the results seen with locomotion (Figure 1). Collectively, these experiments confirm a gene-environment interaction between nicotine and *SV2*, which is statistically significant when quantified with a 2-way ANOVA (p <0.0001 for interaction).

**Figure 2.**
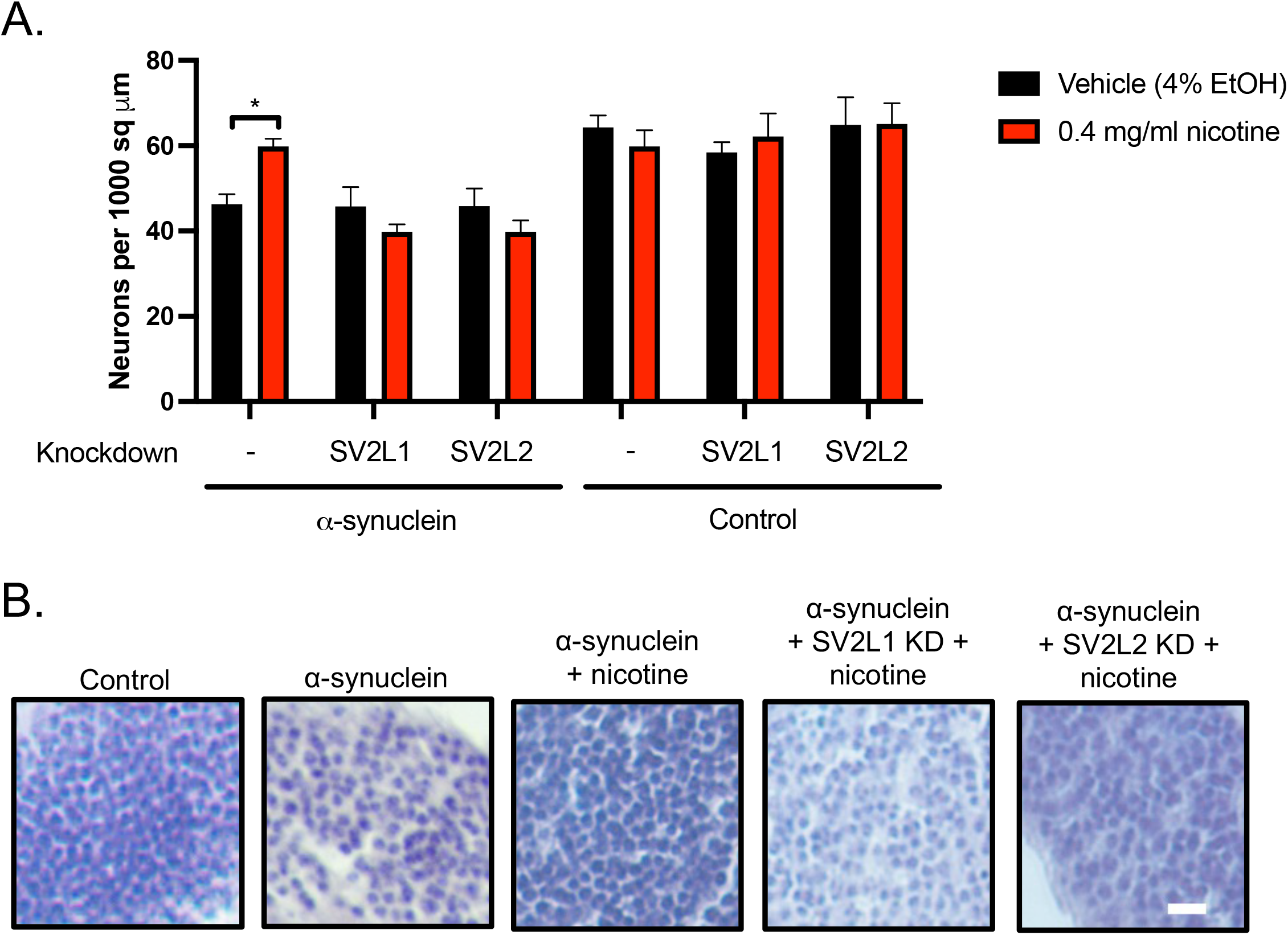
Rescue of neurodegeneration by high dose nicotine requires *SV2*. A. Quantification of total neurons from hematoxylin-stained slides of anterior medulla, n = 6 replicates per genotype. * = p <0.05, ** = p <0.01, determined with one-way ANOVA with Dunnet’s test for multiple comparisons. B. Representative sections of the anterior medulla. Scale = 10 μm.

**Figure 3.**
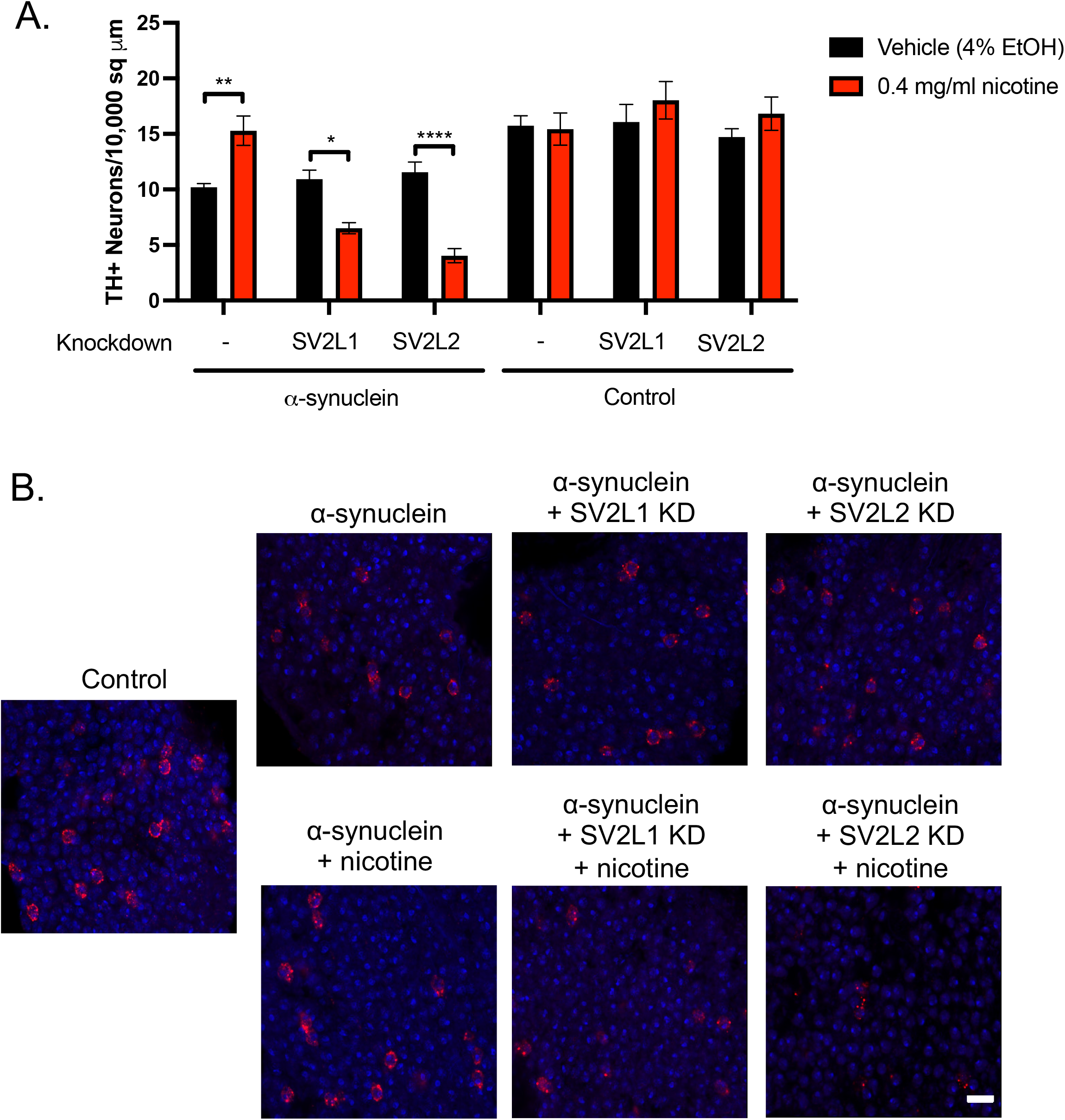
SV2 knockdown and dopaminergic neurons. A. Quantification of dopaminergic neurons. n = 6 replicates per genotype. * = p <0.05, ** = p <0.01, determined with one-way ANOVA with Dunnet’s test for multiple comparisons. B. Anterior medulla sections were stained with tyrosine hydroxylase (red) and mounted with media containing DAPI (blue). Scale = 5 μm.

### Rescue of α-synuclein proteostasis by nicotine requires SV2

We next examined the effect of nicotine on α-synuclein proteostasis. Nicotine reduced the number of large α-synuclein aggregates (Figure 4A, 4C) as well as the number of high molecular weight oligomers (Figure 4B, 4D), but failed to reduce these α-synuclein species when *SV2L1* and *SV2L2* were knocked down. This experiment suggests that the neuroprotective effects of nicotine involve partial normalization of α-synuclein proteostasis, and that this requires *SV2*.

**Figure 4.**
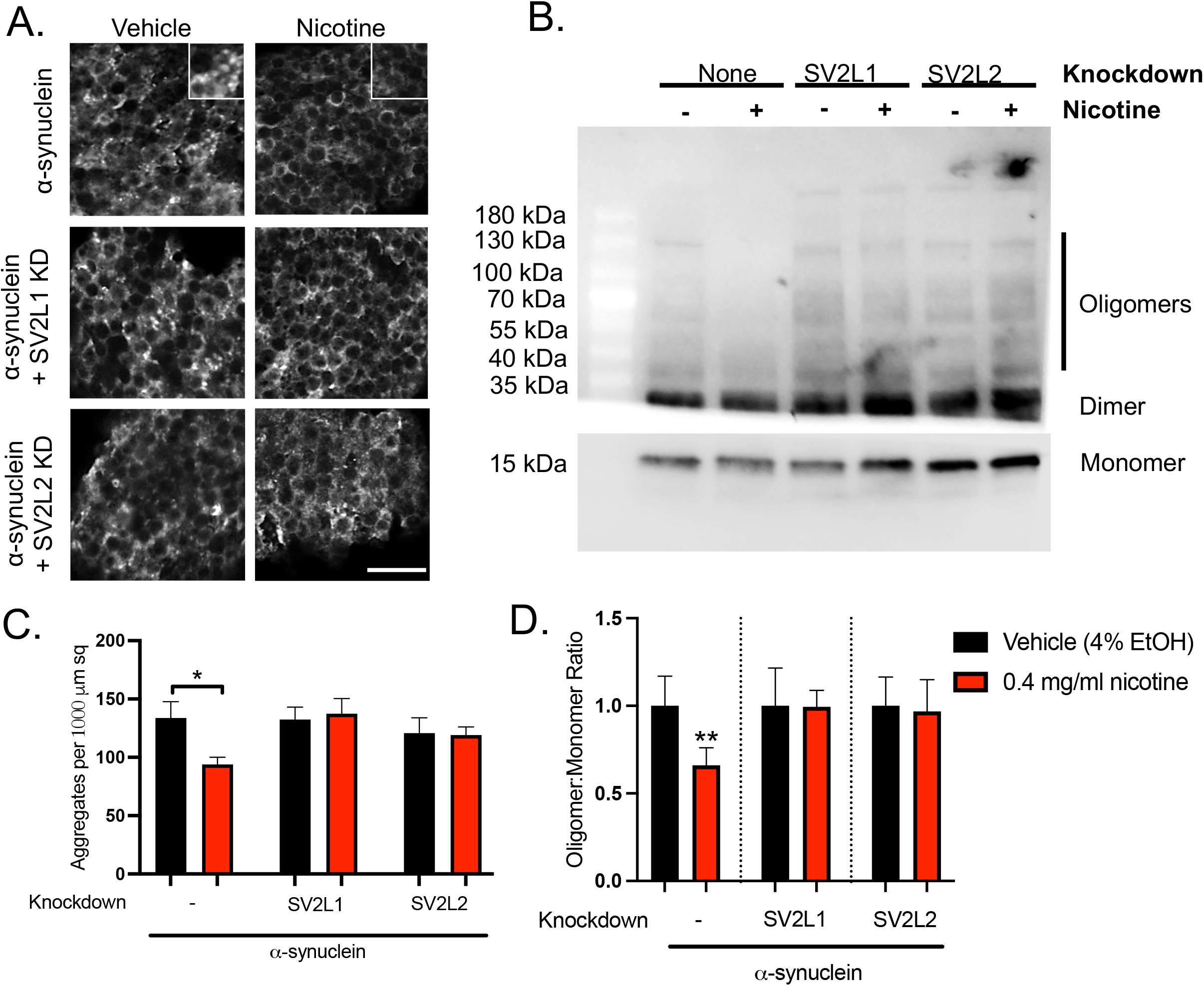
High dose nicotine decreases α-synuclein aggregation and oligomerization. **A.** Slides were stained with anti-α-synuclein (clone 5G4, mouse, 1:50,000) and DAPI. Representative images are shown. **B.** Representative immunoblot demonstrating high molecular oligomers as well as monomeric α-synuclein. **C.** Quantification of α-synuclein aggregates. **D.** Quantification from 4 independent experiments. Each nicotine condition was normalized to its vehicle control, and statistical significance was determined using a 1 sample t-test.

### Nicotine normalizes markers of autophagy

To further investigate the mechanism of the nicotine-*SV2C* interaction on proteostasis, we turned to the lysosomal autophagy system. We have previously investigated the relationship between α-synuclein and autophagy in detail in this model system, finding that α-synuclein expressing flies accumulate autophagosomes due to impaired autophagic flux^27^. Here we examined autophagy using two markers: ref(2)P (the *Drosophila* ortholog of p62) and Atg8a (the *Drosophila* ortholog of LC3). As we have demonstrated previously, α-synuclein transgenic flies demonstrated increased number of puncta staining for both of these markers, as compared to control flies (Figure 5). Further, high-dose nicotine partially rescued this, but failed to do so when *SV2L1* and *SV2L2* was knocked down. This suggests that the neuroprotective effects of nicotine involve the lysosomal autophagy system.

**Figure 5:**
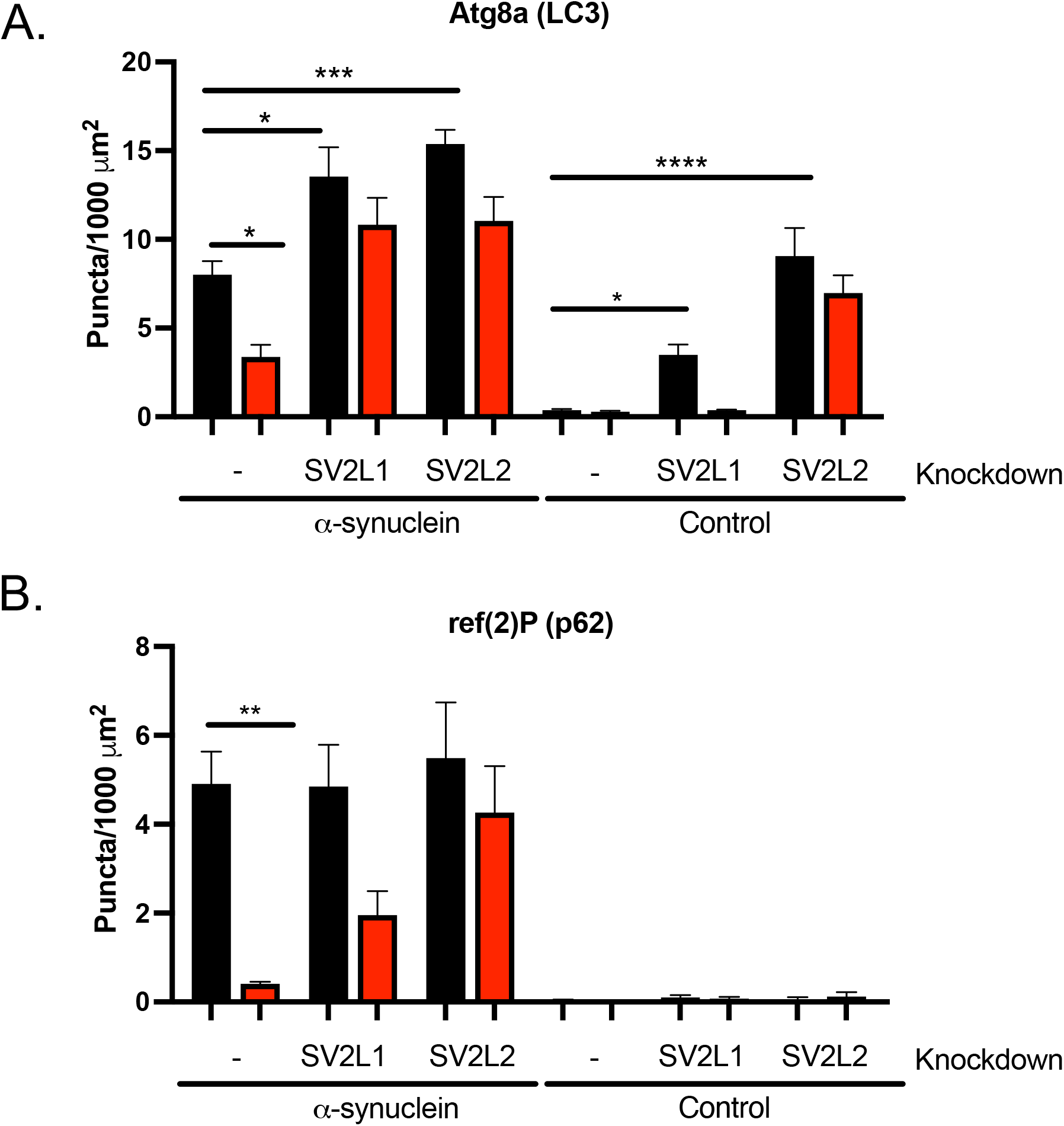
High dose nicotine reduces markers of pathologic autophagy. Immunohistochemistry was performed for ref(2)P (p62), ubiquitin, and Atg8a (LC3). Images were thresholded in ImageJ, and puncta >0.25 μm in area were counted using the Analyze Particles function. n = 6 flies per genotype. **A.** Quantification of ref(2)P (p62) puncta. **B.** Quantification of ubiquitin puncta. **C.** Quantification of Atg8a (LC3 puncta). Knockdown of *SV2* increases Atg8a puncta even in the absence of α-synuclein but has no effect of ref(2)P or ubiquitin puncta. Statistical analysis was performed with a one-way ANOVA with Dunnet’s multiple comparison test. Nicotine and/or *SV2* knockdown conditions were compared to α-synuclein or control flies treated with vehicle and with no RNAi. * = p <0.05, ** = p <0.01, *** = p <0.005, **** = p <0.001.

### SV2C has independent effects on autophagy

Surprisingly, *SV2L1* and *SV2L2* knockdown independently increased accumulation of Atg8a+ puncta (Figure 5A), even in the absence of α-synuclein, though this was not the case for ref(2)P+ puncta (Figure 5B). To confirm the increase in Atg8a+ puncta, we expressed a GFP-tagged Atg8a. As expected, α-synuclein expressing flies demonstrated increased Atg8a-GFP puncta, which was rescued by high-dose nicotine (Supplemental Figure 3A-B). Further, *SV2L1* and *SV2L2* knockdown also increased the number of Atg8a-GFP puncta, even in the absence of α-synuclein (Supplemental Figure 3C-D). Thus, *SV2C* knockdown increases Atg8a+ puncta but not ref(2)P+ puncta.

Atg8a is the *Drosophila* ortholog of LC3, the canonical marker of autophagosomes, and is thus a general marker of autophagosomes. In contrast, ref(2)P, the *Drosophila* ortholog of p62, is found in protein aggregates in the setting of disrupted autophagy, as occurs in neurodegeneration. We therefore hypothesized that *SV2C* knockdown was increasing basal autophagy, independently of an effect on the pathologic autophagosomes that accumulate with α-synuclein. To investigate this hypothesis, we used a tandem tag reporter, GFP-mCherry-Atg8a. This reporter expresses both GFP and mCherry in newly formed autophagosomes, but the GFP fluorescence is quenched in the acidic autolysosome. Thus, a large mCherry to GFP ratio indicates preserved autophagic flux, whereas a small mCherry to GFP ratio indicates impaired autophagic flux.

We have previously reported that α-synuclein expressing flies have impaired autophagic flux using this reporter, as indicated by many dual GFP and mCherry positive puncta (Sarkar, 2021). Here we again demonstrate that α-synuclein expression leads to impaired autophagic flux, with more dual-positive puncta and a low mCherry to GFP ratio compared to control flies (Figure 6A-B, 6F). The mCherry to GFP ratio was normalized by either treatment with nicotine or by *SV2C* knockdown (Figure 6C-D, 6F). However, when both SV2C knockdown and nicotine treatment were present, autophagic flux was not normalized (Figure 6E-F). Thus, nicotine increases autophagic flux in α-synuclein expressing flies in an SV2C-dependent manner. In the absence of α-synuclein, *SV2C* knockdown led to an increase in the mCherry to GFP ratio (Figure 6G), supporting the idea that *SV2C* knockdown is increasing basal autophagy. Treatment with nicotine blunted this effect (Figure 6G).

**Figure 6:**
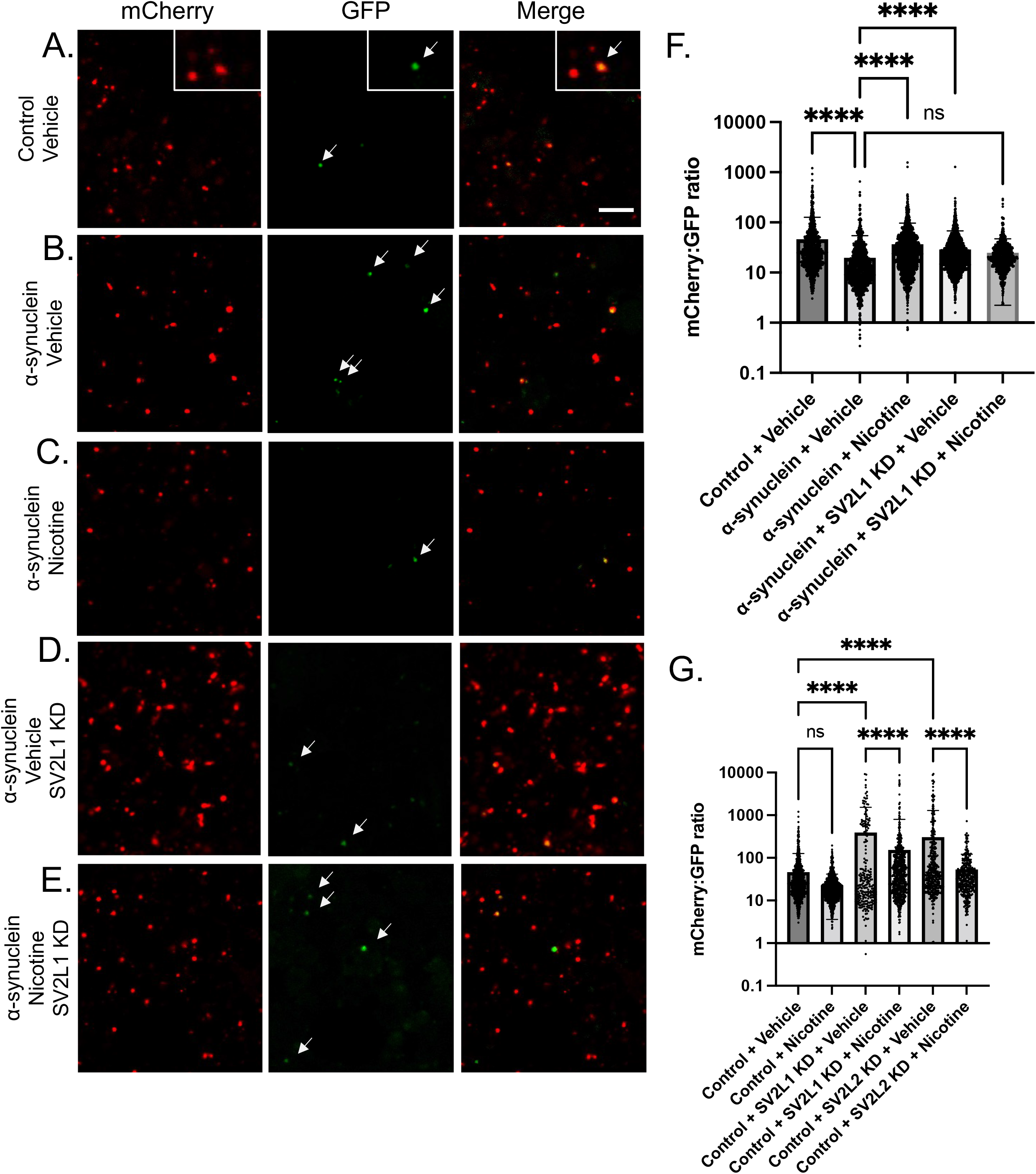
SV2 knockdown increases physiologic autophagy. A. *UAS-GFP-mCherry-Atg8a* was expressed in control or α-synuclein flies, with or without *SV2L2* knockdown, treated with nicotine or vehicle control. Brains were dissected in PBS and imaged immediately and sequentially in the red and green channels. A. In control flies, numerous mCherry+ puncta are seen, with exceedingly rare dual positive puncta (see inset). A plot profile of red and green fluorescence was calculated in ImageJ and is shown for two representative puncta. The left plot corresponds to the left punctum, and the right plot corresponds to the right punctum. Scale = 5 μm. B. Images. Scale is the same as in (A). C. Fluorescence intensity of mCherry:GFP was averaged across all puncta. n = 3 flies per condition. Genotype is *UAS-GFP-mCherry-Atg8a/+; nSybQF2, Snyb-GAL4/UAS-CG14691 RNAi*.

## Discussion

Here, we demonstrate that nicotine rescues neurodegeneration in a genetic model of PD in which human α-synuclein is over-expressed in *Drosophila* neurons. While multiple previous investigations have identified a neuroprotective effect of nicotine in toxic models of PD, including MPTP^1,11^, 6-OHDA^29^, rotenone^13^, and paraquat^2^, studies involving genetic models are more limited. Nicotine has been examined in a *parkin* mutant *Drosophila* model^14^, as well as recent paper involving overexpression of *Synphilin-1* in dopaminergic neurons, also in *Drosophila*^30^. Surprisingly, few studies have examined nicotine in α-synuclein based models of PD. Bono et. al found a protective effect of nicotine on α-synuclein accumulation in cultured mouse and human DA neurons^31^, and Kardani et. al found that nicotine slowed α-synuclein oligomerization in yeast^32^. To our knowledge, there is only one published study involving nicotine in α-synuclein-based PD model in a multicellular organism *in vivo:* Subramaniam and colleagues investigated nicotine in α-synuclein over-expressing transgenic mice, finding improvement in cognitive performance no improvement in neurodegeneration, α-synuclein aggregation, or motor impairment^33^.

Beyond investigating a role for nicotine in an α-synuclein-based model of PD, we sought to determine the mechanism of interaction between nicotine and *SV2*. Of interest, both *SV2C* and nicotine have been reported to affect release of DA. Using fast scan cycle voltammetry, Dunn et. al determined that nicotine increases DA release in response to high-intensity stimulation in the dorsal striatum of wild-type mice, but when tested in *SV2C* knockout animals, nicotine paired with high-intensity stimulation reduced DA release to less than half of the response at baseline^7^. Thus, SV2C levels determined the response of dopaminergic neurons to nicotine. In line with this, we found that nicotine rescued motor functioning and prevented DA neuron death in α-synuclein flies, but when *SV2* orthologs were knocked down, nicotine not only failed to rescue but in fact significantly further decreased locomotion and DA neuron survival. Together, these results suggest that SV2 expression level mediates the effects of nicotine on DA neurons, ranging from protective to harmful.

Interestingly, beyond the effects on DA neurons, we found that the nicotine x *SV2* interaction also influenced α-synuclein proteostasis. That is, nicotine treatment rescued α-synuclein oligomerization and aggregation in the presence of wildtype *SV2L1* and *SV2L2* expression, but it failed to do so in *SV2L1* or *SV2L2* knockdown conditions (Figure 4). Of interest, Dunn et al reported a direct protein-protein interaction between SV2C and α-synuclein, an increase in high molecular weight α-synuclein in *SV2C* knockout mice, and an abnormal *SV2C* expression pattern in an A53T α-synuclein mouse model and PD patient brains. How nicotine might further influence any interaction between α-synuclein and *SV2C* is unknown.

To further understand the effects of the nicotine x *SV2* interaction on proteostasis, we examined autophagy. Autophagy is a highly regulated process – it is necessary for cellular homeostasis and response to stress, but when excessive, it causes apoptosis^34^. Further, there is a complex relation between autophagy and α-synuclein. Alterations in the lysosomal autophagy system increase α-synuclein accumulation, and excessive and/or abnormal α-synuclein then further impedes the lysosomal autophagy system in a vicious cycle^35^. Of note, nicotine has been shown to induce autophagy in several tissues outside of the brain, though the effects of this range from beneficial to harmful depending on the tissue^36–43^. Here we found that nicotine treatment rescued impaired autophagic flux in α-synuclein flies, and that this rescue was dependent on intact expression of *SV2C* (Figure 6-7).

**Figure 7:**
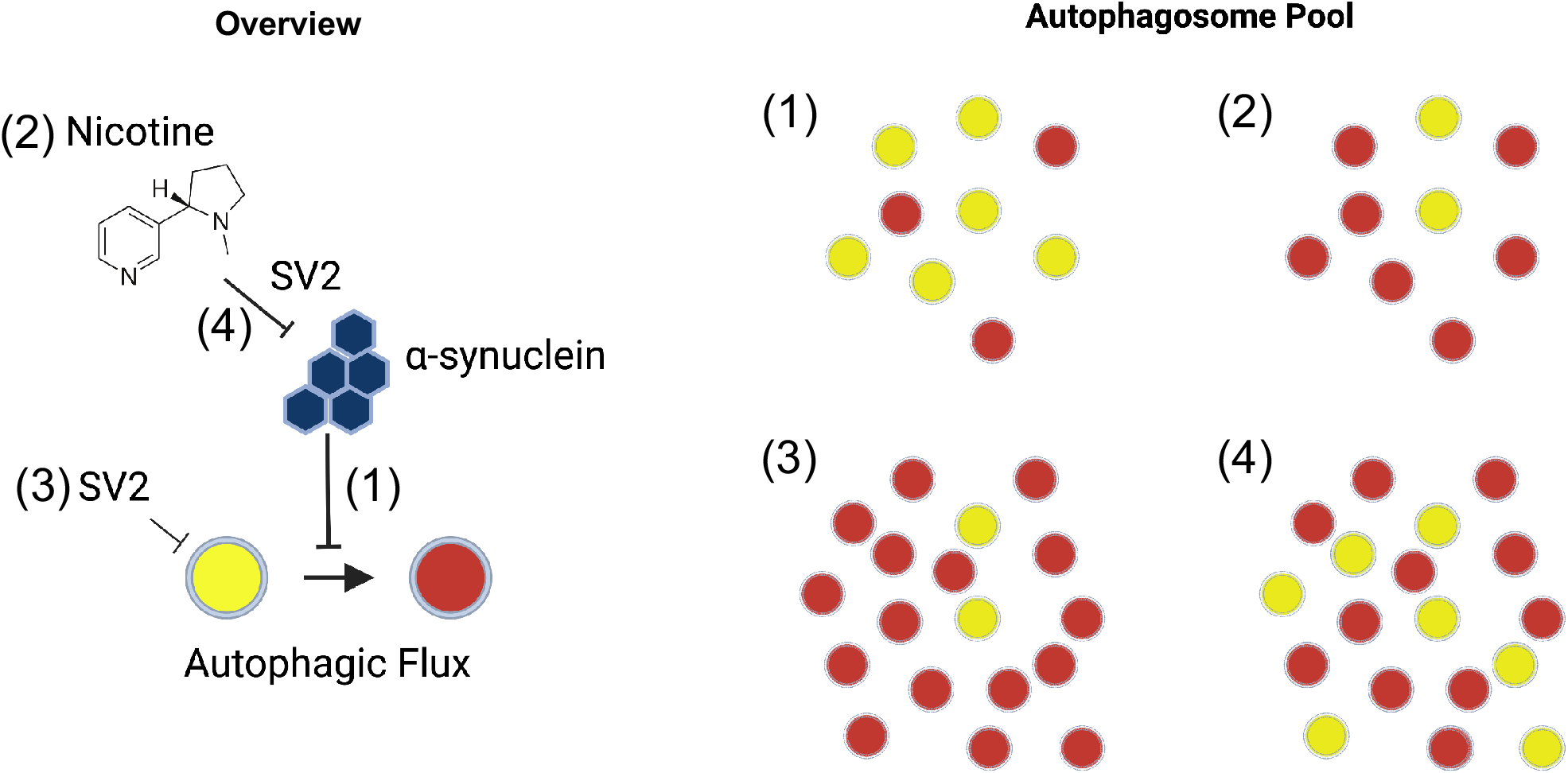
Proposed model. 1. α-synuclein expressing flies have impaired autophagic flux. 2. Nicotine can restore autophagic flux in α-synuclein flies in an SV2-dependent manner. 3. *SV2* knockdown increases basal autophagy, resulting in more autophagosomes but no delay in maturation. 4. In the absence of SV2C nicotine is unable to rescue autophagic flux, perhaps because the increase in basal autophagy resulting in more autophagosomes overwhelms the autophagy-lysosomal system.

Unexpectedly, we also found an effect of *SV2C* on autophagy that was independent of α-synuclein. That is, knockdown of *SV2L1* or *SV2L2* in either control or α-synuclein animals increased the number of autophagosomes, as indicated by the canonical autophagy marker Atg8a (LC3) (Figure 5A). We believe that this is due most likely to an increase in basal autophagy levels, as ref(2)P (p62) levels were not increased by *SV2L1* or *SV2L2* knockdown (Figure 5B), and *SV2L1* or *SV2L2* knockdown led to an increase in autophagic flux (Figure 6-7). How SV2C may regulate autophagy is unknown, but it is worth mentioning that SV2 family proteins are transmembrane proteins that are thought to be present on all secretory vesicles, and they play a role in vesicular transport, trafficking, anchoring, and recycling^44^. If this membrane interaction affects other membranous organelles, like autophagosomes or lysosomes, this could explain the varied effects of SV2 observed in our study.

In summary, we show here that nicotine-mediated rescue of α-synuclein toxicity requires SV2. This work validates the gene-environment interaction between nicotine and *SV2* and provides insight on the mechanism, finding an unexpected role for this interaction in α-synuclein proteostasis and autophagy. Further, the work provides proof of concept of the utility of this model for investigating additional GxE interactions. Understanding these GxE is critical for future clinical trial design and identifying which patients may be most likely to benefit from which therapies.

## Acknowledgement

The HC3 α-synuclein antibody and GFP antibodies were provided by the Developmental Studies Hybridoma Bank, created by the NICHD of the NIH and maintained at The University of Iowa, Department of Biology, Iowa City, IA 52242. *Drosophila* stocks obtained from the Bloomington *Drosophila* Stock Center (NIH P40OD018537) were used in this study. We thank the Transgenic RNAi Project (TRiP) at the Harvard Medical School (NIH-NIGMS R01GM084947) for making transgenic RNAi stocks.

## Authors’ Roles

ALO data acquisition, data analysis and interpretation, manuscript writing and revision. SGC data acquisition, manuscript writing. MBF acquisition of funding, project design, data interpretation, and manuscript revision.

## Financial Disclosures for all authors

MBF receives grant support from NIH-NINDS, NIH-NIA, and the Michael J Fox Association and Aligning Science Across Parkinson’s (ASAP) initiative. ALO receives grant support NIH-NINDS, the Department of Defense, and the American Parkinson’s Disease Association. SGC has no financial disclosures.

## Supplemental Figure Legends

**Supplemental Figure 1**. **Confirmation of locomotion rescue with a second RNAi line. α-** synuclein or control flies with or without knockdown of *SV2L1* and *SV2L2* using VDRC lines. Flies were aged to 7 days post-eclosion on food containing 0.4 mg/ml nicotine or vehicle control. Symbols above the 0.4 mg/ml nicotine bar represent statistically significant difference compared to the vehicle control, respectively. *p <0.05, **p < 0.01, ***P<0.005, ns = p > 0.05. n = minimum of 60 flies per genotype (6 biological replicates of 10 flies each). Significance was determined by two-way ANOVA with Sidak’s test for multiple comparisons.

**Supplemental Figure 2. Confirmation of knockdown.** A. qRT-PCR for *SV2L1* in control flies (no RNAi) or flies expressing one of two RNAi’s targeting *SV2L1*. RNAi #1 is the JF line. RNA #2 is the VDRC line. B. qRT-PCR for *SV2L2* in control flies (no RNAi) or flies expressing one of two RNAi’s targeting *SV2L2*. RNAi #1 is the JF line. RNA #2 is the VDRC line.

**Supplemental Figure 3. *SV2* knockdown increases Atg8a even in the absence of α-synuclein**. **A.** Representative images. **B.** Quantification. Images were thresholded in ImageJ, and puncta > 0.12 μm in area were counted using the Analyze Particles function. Control flies expressing a GFP tagged Atg8a treated with ethanol vehicle or nicotine demonstrate sparse small puncta (small white arrows). Puncta are larger and more numerous with α-synuclein and large puncta are reduced in number with nicotine treatment (large yellow arrows). Scale = 5 μm. n = 6 flies per condition. Genotypes are *nSyb-QF2, nSyb-GAL4/UAS-Atg8a-GFP* or *QUAS-Syn, nSyb-QF2, nSyb-GAL4/UAS-Atg8a-GFP* for control and α-synuclein expressing flies, respectively. **C.** Representative images. **D.** Quantification of puncta was performed as in B. Genotypes are *nSyb-QF2, nSyb-GAL4, UAS-Atg8a-GFP/+, nSyb-QF2, nSyb-GAL4, UAS-Atg8a-GFP/UAS-CG3168 RNAi* or *nSyb-QF2, nSyb-GAL4, UAS-Atg8a-GFP/UAS-CG14691 RNAi*.

## Notes

**Funding Sources**, This work was funded by 1R21 NS0105151 (MBF), 1R01 NS098821 (MBF), 5K08-K08NS109344-03 (ALO), and W81XWH-18-1-0395 (ALO). The study is funded by the joint efforts of The Michael J. Fox Foundation for Parkinson’s Research (MJFF) and the Aligning Science Across Parkinson’s (ASAP) initiative. MJFF administers the grant [ASAP-000301] on behalf of ASAP and itself.

### Competing Interest Statement

The authors have declared no competing interest.

