## Supplemental Figures for "Nicotine-mediated rescue of α-synuclein toxicity requires synaptic vesicle glycoprotein 2"

### Slide 1
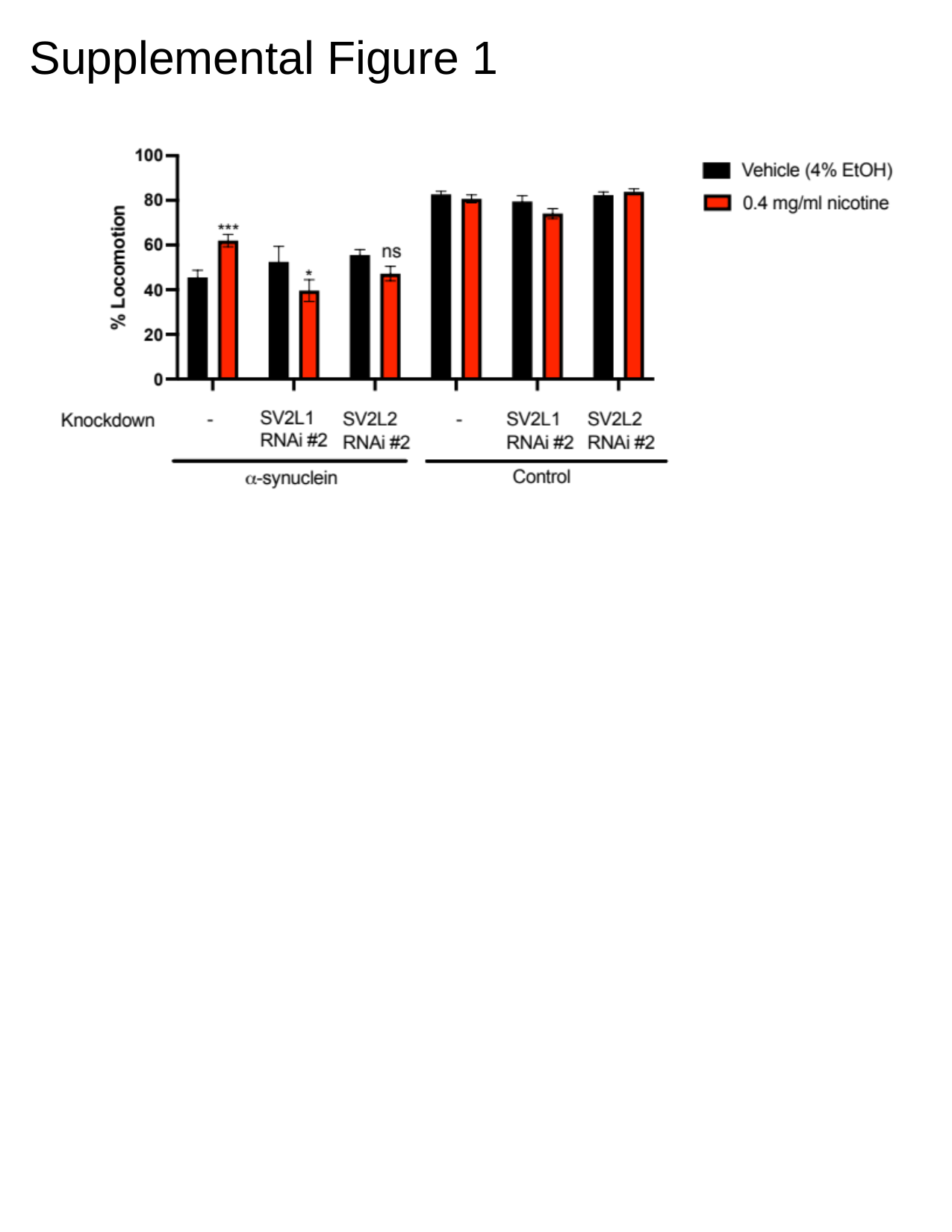

#### Supplemental Figure 1

### Slide 2
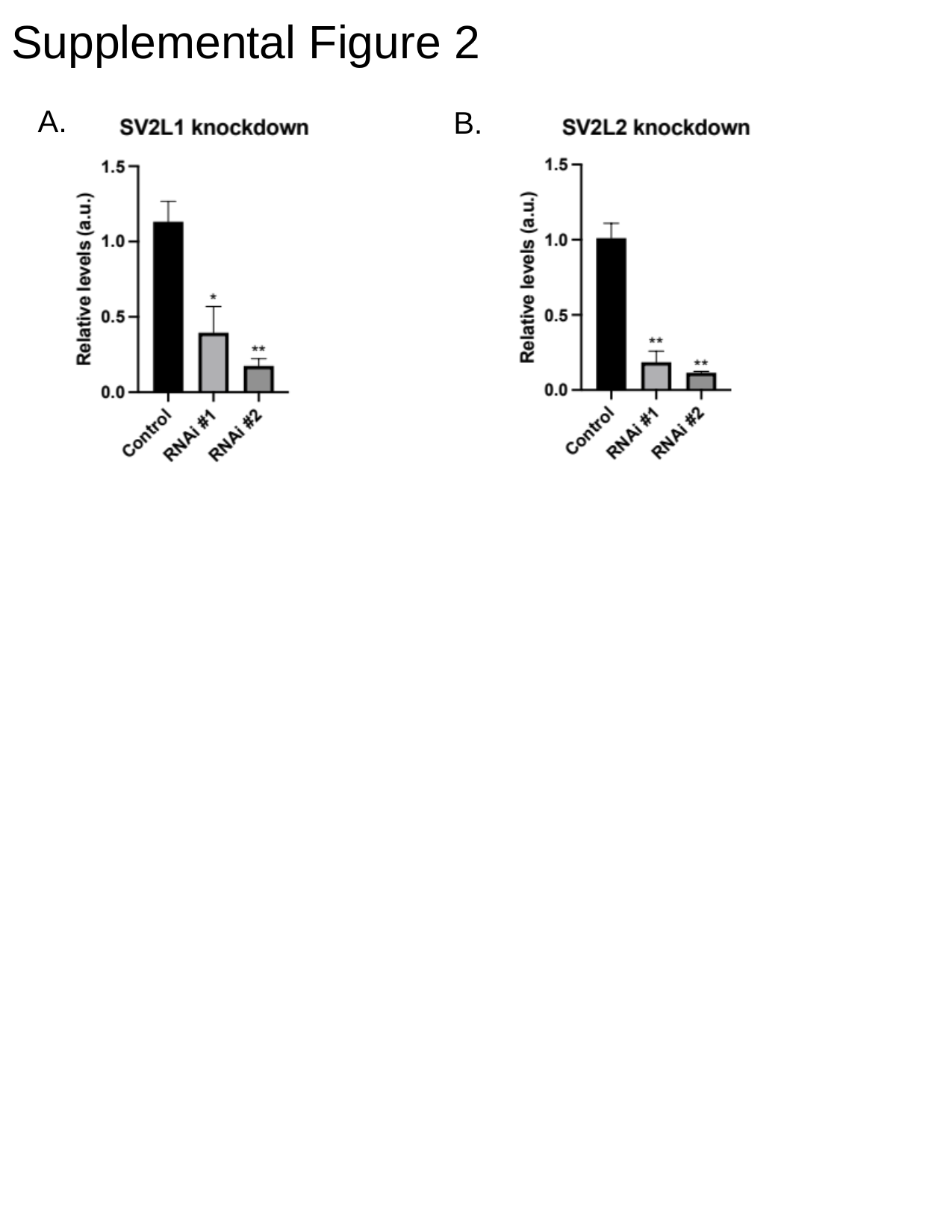

#### Supplemental Figure 2
A.
B.

### Slide 3
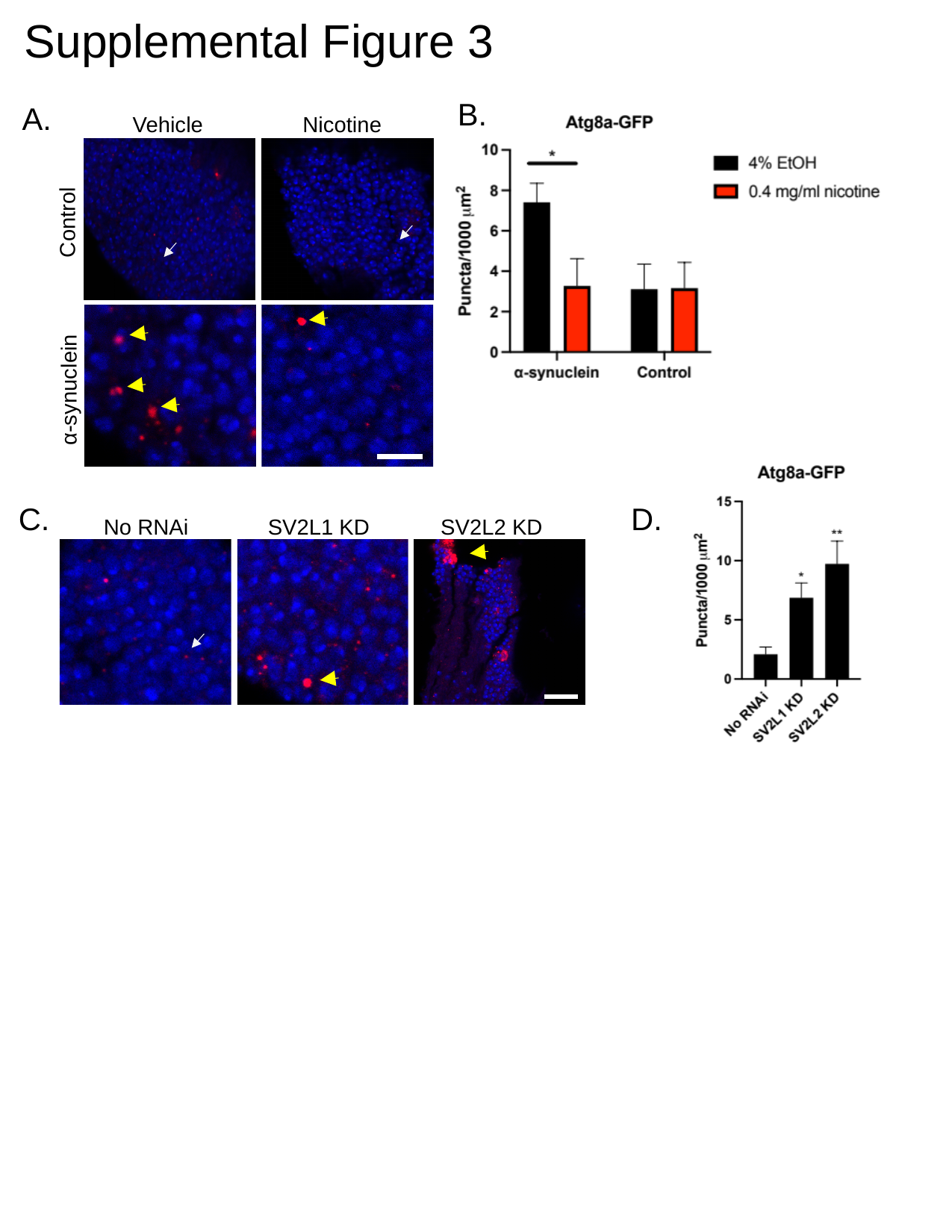

Supplemental Figure 3
B.
A.
Nicotine
Vehicle
Control
α-synuclein
C.
D.
SV2L1 KD
SV2L2 KD
No RNAi
